# RNase L activating 2′–5′ oligoadenylates bind ABCF1, -3 and Decr-1

**DOI:** 10.1101/2023.03.21.532770

**Authors:** Apurva A. Govande, Aleksandra W. Babnis, Christian Urban, Matthias Habjan, Rune Hartmann, Philip J. Kranzusch, Andreas Pichlmair

## Abstract

A notable signaling mechanism employed by mammalian innate immune signaling pathways uses nucleotide based second messengers such as 2′–3′-cGAMP and 2′–5′-oligoadenylates (2′–5′ OA), which bind and activate STING and RNase L, respectively. Interestingly, the involvement of nucleotide second messengers to activate antiviral responses is evolutionary conserved, evidenced by the identification of an antiviral cGAMP-dependent pathway in *Drosophila*. Using a mass spectrometry approach, we identified several members of the ABCF family in human, mouse, and *Drosophila* cell lysates as 2′–5′ OA binding proteins, suggesting an evolutionary conserved function. Biochemical characterization of these interactions demonstrates high-affinity binding of 2′–5′ OA to ABCF1, which depended on phosphorylated 2′–5′ OA and an intact Walker A/B motif of the ABC cassette of ABCF1. As further support for species-specific interactions with 2′–5′ OA, we additionally identified that the metabolism enzyme Decr1 from mouse, but not human or *Drosophila* cells forms a high-affinity complex with 2′–5′ OA. A 1.4 Å co-crystal structure of the mouse Decr1–2′–5′ OA complex explains high-affinity recognition of 2′–5′ OA and the mechanism of species-specificity. Despite clear evidence of physical interactions, we could not identify profound antiviral functions of ABCF1, ABCF3 or Decr1 or 2′–5′ OA-dependent regulation of cellular translation rates as suggested by the engagement of ABCF proteins. Thus, although the biological consequences of the here identified interactions need to be identified, our data suggests that 2′–5′ OA can serve as signaling hub to distribute a signal to different recipient proteins.

## Importance

Small nucleotides can serve as essential components of the innate immune system signaling system. Here we report that the 2’-5’ OA can bind to a wider repertoire of proteins than anticipated. In particular, we identified proteins of the ABCF family as well as murine Decr1 as high affinity binders and characterize these interactions using biochemical experiments and crystallography. Moreover, we tested a wide array of biological and virological functions to evaluate the involvement of the identified proteins in antiviral immunity.

## Introduction

Oligoadenylate synthase (OAS) proteins are cytosolic double-stranded (dsRNA) sensors that detect viral infection and enzymatically synthesize the nucleotide second messenger 2′–5′ oligoadenylate (2′–5′ OA) (1–5). Humans encode three catalytically active OAS enzymes, OAS1, OAS2, and OAS3, that vary in size and subcellular localization (6), and OASL, a catalytically inactive OAS-like protein that contains two ubiquitin-like domains and has been implicated in regulating RIG-I activity (7–9). For all three active OAS proteins, sensing of dsRNA leads to a conformation change whereupon the catalytic domain synthesizes the linear signaling molecule 2′–5′ OA composed of adenine nucleotides linked through noncanonical 2′–5′ phosphodiester bonds (10). In human cells, OAS1, OAS2, and OAS3 have been shown to be involved in the antiviral innate immune response against a wide range of viruses with OAS3 being particularly important for viral dsRNA detection (11–14). The product 2′–5′ OA molecules vary in size between 3-and 30-nucleotide long chains (15). Chains between three to five bases of 2′–5′ OA bind and activate a latent endoribonuclease, RNase L, in the cytosol of cells (3, 16). RNase L is known to bind 2′–5′ OA with ∼10 nM affinity, and high affinity RNase L–2′–5′ OA interaction requires specific recognition of the 2′–5′ OA 5′ triphosphate (17, 18). Upon activation, RNase L promiscuously cleaves single stranded RNA, resulting in degradation of host and viral mRNA transcripts and cellular ribosomal RNA. RNase L activation leads to translational arrest and apoptosis, thereby establishing a broadly antiviral state in the cell (19, 20). RNase L is not activated by 2′–5′ OA shorter than three base pairs in length (15), and there is no defined role of 2′–5′ OA for chains longer than 5 bases.

OAS proteins are evolutionarily ancient and widely conserved in metazoans. In contrast, RNase L is limited to jawed vertebrates and is surprisingly absent in invertebrates and primitive metazoan lineages (21). In mice, OAS proteins have undergone recent duplication events, leading to an expanded panel of OAS functions and OAS-like proteins (22–24), including OAS2, OAS3, OASL, and eight paralogs of OAS1 (25). Evidence of evolutionary pressure in the OAS family, such as duplication and diversification events (25), positive selection at key nucleic acid binding regions (26), as well as the absence of RNase L in more ancient organisms suggests a wider role for 2′–5′ OA signaling independent of RNase L. Here, we interrogated RNase L-independent roles of 2′–5′ OA using an unbiased approach to identify alternative 2′–5′ OA binding partners by affinity purification and high-resolution mass spectrometry. Our results identified novel 2′–5′ OA associating proteins, which suggest a role of 2′–5′ OA in signal distribution and that may point towards auxiliary roles of OAS-signaling pathways.

## Results and Discussion

### Identification of 2′–5′ OA binding proteins in eukaryotic cells

To identify proteins that associate with OAS products, we enzymatically generated biotinylated 2′–5′ OA using human OAS1 reactions that produce chains between 3–10 nucleotides (9, 15, 27). 2′–5′ OA or ATP were coupled to beads and used for affinity purification from human, mouse, or *Drosophila melanogaster* cell lysates followed by mass spectrometry-based protein identification (Figure 1a). As expected, 2′–5′ OA but not ATP precipitated RNase L from human and mouse cell lysates, confirming validity and specificity of this approach (Figure 1b-c). Notably, besides RNase L we found a highly significant enrichment of proteins belonging to the ABC superfamily, with the highest enrichment for proteins of the ABCF family including ABCF1, -3 in human and mouse, as well as ABCF2 in mouse (Figure 1b-c; Supplementary Table S1). An additional identified 2′–5′ OA interactor was murine Decr1, which is a mitochondria-resident protein involved in beta-oxidation and metabolism of unsaturated fatty enoyl-CoA esters. Remarkably, flies, that do not express RNase L, similarly enriched for CG1703, CG9330-RA and pix, which are orthologues of ABCF1, -3 and the closely related protein ABCE1, respectively (Figure 1d; Supplementary Table S1). The identification of the same proteins across distantly related species suggests an evolutionary conservation of these interactions. ABCF1 (also known as ABC50) was first identified as a protein binding to eIF2 (28), and is involved in translation initiation and elongation (29). Abcf1 deficient mice are embryonically lethal, which may be explained by the involvement of this protein in translation (30). Intriguingly, ABCF1 has also been proposed to have immune-regulatory functions due to binding to DNA and the ability to regulate type-I IFN induction in response to cytosolic delivery of DNA (31). Similarly, murine Abcf3 was shown to exert an antiviral activity against flaviviruses (23).

**Figure 1.**
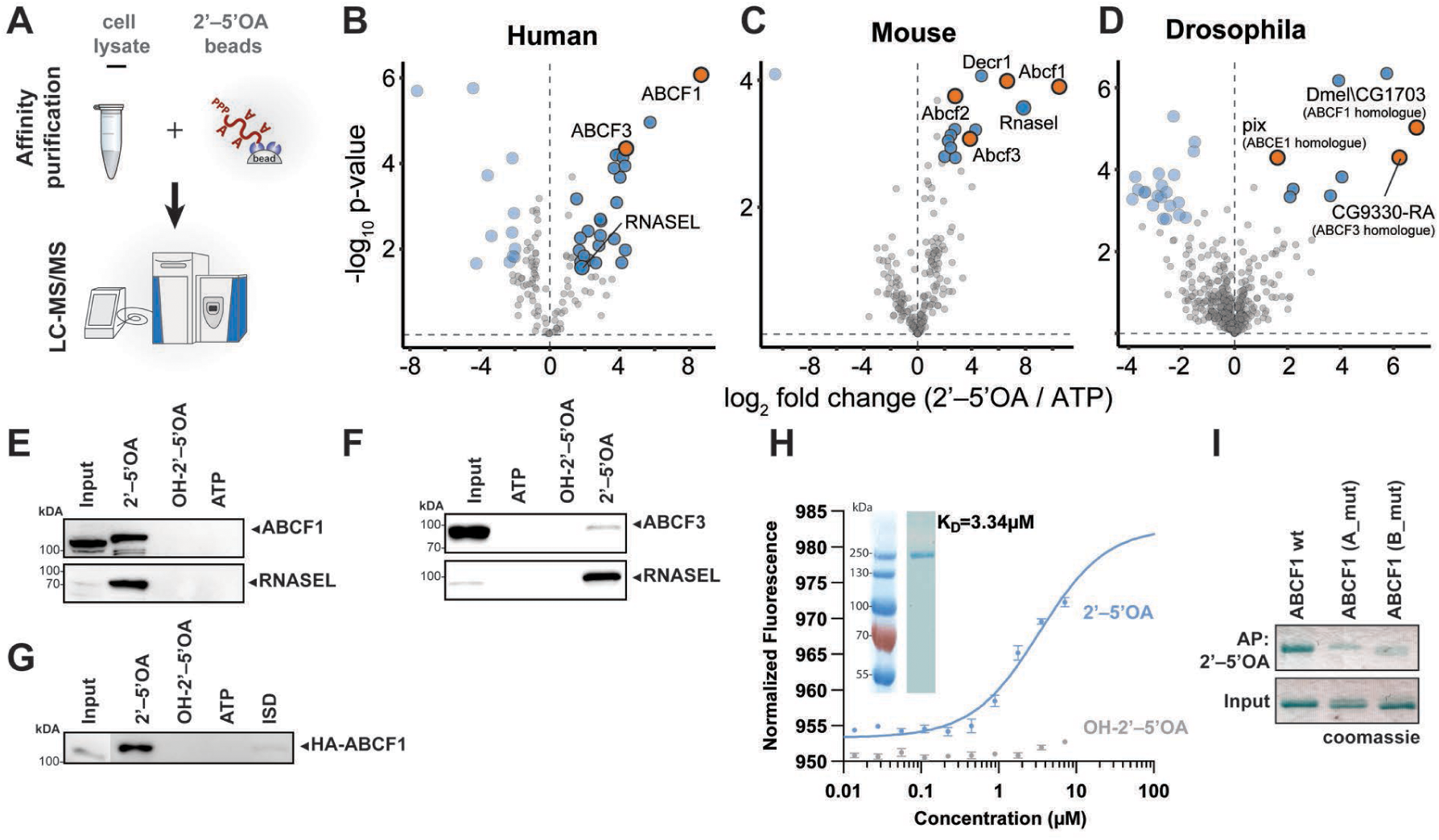
Identification of 2′–5′ OA binding proteins using affinity purification mass spectrometry. **(A)** Scheme of affinity purification with 2′–5′ OA conjugated beads and human, fly and mouse lysates for mass spectrometry. Cell lysates were incubated with beads linked to ATP or 2′–5′ OA, followed by affinity enrichment and analysis of bead-bound proteins using LC-MS/MS. **(B-D)** Identification of 2′–5′ OA binding proteins in **(A)** human (HeLa S3, n = 4), **(B)** mouse (BMDMs, n = 3) and **(D)** *Drosophila* (Schneider S2, n = 4) cell lysates. Volcano plots show the log_2_ fold change enrichment of proteins in 2’–5’ OA vs ATP samples (x-axis) plotted against the log_10_ transformed p-value (y-axis). Proteins in blue are significantly enriched (two-sided Student’s t-test, S0 = 0, FDR < 0.01, log_2_ fold change ≥ 1.5) and proteins in orange are significantly enriched hits belonging to the ABCE and ABCF subfamilies/ABC superfamily. **(E, F)** Validation of ABCF1, -3 and RNAse L binding to the indicated affinity beads in HeLa (E) and THP-I (F) cell lysates that were treated with type-I IFN overnight. **(G)** as for (E) but 293T cell lysates from HA-ABCF1 overexpressing cells were used. **(H)** Binding affinity of recombinant ABCF1 to fluorescent 2′–5′ OA or OH-2′–5′ OA as determined by microscale thermophoresis (MST). The inset shows a coomassie stain on a SDS-PAGE of the recombinant protein. The graph shows mean ± sd for 2′–5′ OA (n = 4) and OH-2′–5′ OA (n = 3). **(I)** Precipitation of recombinant ABCF1 and the indicated variants of ABCF1 with mutations in walker A and B motifs with 2′–5′ OA and visualization by coomassie stain.

We confirmed binding of endogenous ABCF1 and ABCF3 to 2′–5′ OA in HeLa and THP-1 cells (Figure 1e, g). The association of RNase L with 2′–5′ OA requires a triphosphate group on the 5′ terminus. Indeed, dephosphorylation of 2′–5′ OA by phosphatase treatment (designated as OH-2′–5′ OA) abolished RNase L binding, as expected. Similarly, binding of ABCF1 and ABCF3 required full phosphorylation of 2′–5′ OA since neither of these proteins precipitated with OH-2′–5′ OA (Figure 1e-f). The interaction between 2′–5′ OA could also be recapitulated with HA-tagged versions of ABCF1 expressed *in trans* in HEK293T cells (Figure 1g). HA-ABCF1 also precipitated with ISD (31), but this interaction was weaker as compared to interaction of HA-ABCF1 with 2′–5′ OA (Figure 1g). To establish whether the interaction between 2′–5′ OA and ABCF1 is direct, we generated recombinant ABCF1 and measured its affinity to fluorescently labelled 2′–5′ OA using microscale thermophoresis. This assay suggested a K_D_ of ∼3 µM between ABCF1 and 2′–5′ OA. In agreement with precipitation experiments, the binding required a triphosphate group (Figure 1h). The nucleotide binding domains of ABC proteins contain two conserved motifs: Walker A and Walker B. Functionality of both motifs was required for efficient precipitation of recombinant ABCF1 (Figure 1i). Collectively, we identified proteins of the ABCF family as well as murine Decr1 as proteins that, similar to RNase L, associate with 2′–5′ OA. The associations described here required a 5′ terminal triphosphate group and mutational analysis of ABCF1 suggest that ABCF family members require a fully functional nucleotide-binding domain.

### ABCF1 and -3 do not affect RNAseL activity or virus growth in cell lines

We next evaluated the possible antiviral roles of ABCF1 in loss of function experiments in different human HeLa-based cell types that do (HeLa S3) or do not (HeLa) express RNase L (Figure 2a). ABCF1 was successfully depleted by siRNA in HeLa S3 and HeLa cells (Figure 2b, c). As expected, poly-I:C transfection of HeLa S3 cells resulted in characteristic ribosomal RNA (rRNA) cleavage, an indication for RNase L activation (Figure 2d). siRNA-mediated depletion of ABCF1 in HeLa cells did not affect rRNA cleavage in response to poly-I:C transfection, suggesting that ABCF1 does not directly interfere with RNase L activity (Figure 2d). In line with reduced RNase L levels in HeLa cells, poly-I:C transfection did not induce rRNA cleavage and depletion of ABCF1 did not affect rRNA quality (Figure 2e). We next tested growth of encephalomyocarditis virus (EMCV) and LaCrosse virus (LaCV) after poly-I:C or mock stimulation of HeLa S3 and HeLa cells. EMCV growth was dramatically reduced in poly-I:C treated HeLa S3 cells (Figure 2f), which is in line with the reported type-I interferon and RNase L sensitivity of this virus (19). Depletion of ABCF1 did not affect virus growth in the absence or presence of poly-I:C. LaCV did not replicate to high virus titers in HeLa S3 cells and loss of ABCF1 did not rescue LaCV replication (Figure 2g). Collectively, these data indicated that ABCF1 does not contribute, or only mildly contributes, to antiviral immunity in presence of RNase L. A possible reason for the apparent lack of response could be a dominant role of RNase L, which could mask potential effects of ABCF1. We therefore used HeLa cells that do not express RNase L and which do not show rRNA cleavage after poly-I:C transfection (Figure 2a, e). Compared to the effects seen in HeLa S3 cells, inhibition of EMCV growth by poly-I:C transfection was less pronounced (Figure 2h), which may be explained by the lack of RNase L. However, depletion of ABCF1 did not affect growth of EMCV (Figure 2h). Similarly, LaCV generally grew to higher titers in HeLa as compared to HeLa S3 cells but the depletion of ABCF1 did not change growth properties of this virus (Figure 2i). These data indicate that the lack of ABCF1 does not, or only mildly affect growth of EMCV and LaCV in HeLa cells. To establish whether other viruses are sensitive to ABCF1 or ABCF3, we employed a CRISPR/Cas9 approach to deplete both proteins in A549 cells and infected them with different fluorescent reporter viruses (Vaccinia virus, VACV; Herpes Simplex virus 1, HSV-1; Vesicular Stomatitis virus, VSV; Rift Valley Fever virus, RVFV and Yellow Fever virus, YFV) and followed growth of these viruses using time-lapse microscopy. As compared to non-targeting control cells (NEG1, NEG2), growth of most viruses was not increased by depletion of ABCF1 or ABCF3 (Figure 2j). The only exception was a minor increase of VACV replication in absence of ABCF3. In contrast, YFV growth was reduced in the absence of ABCF1, suggesting a pro-viral effect of this protein for flaviviruses. Collectively, our data indicate that ABCF1 and ABCF3 have little or no antiviral effect on growth of the majority of viruses, suggesting that, in the tested human cells, these proteins and their interaction with 2′–5′ OA may have alternative roles that is not directly linked to antiviral immunity.

**Figure 2:**
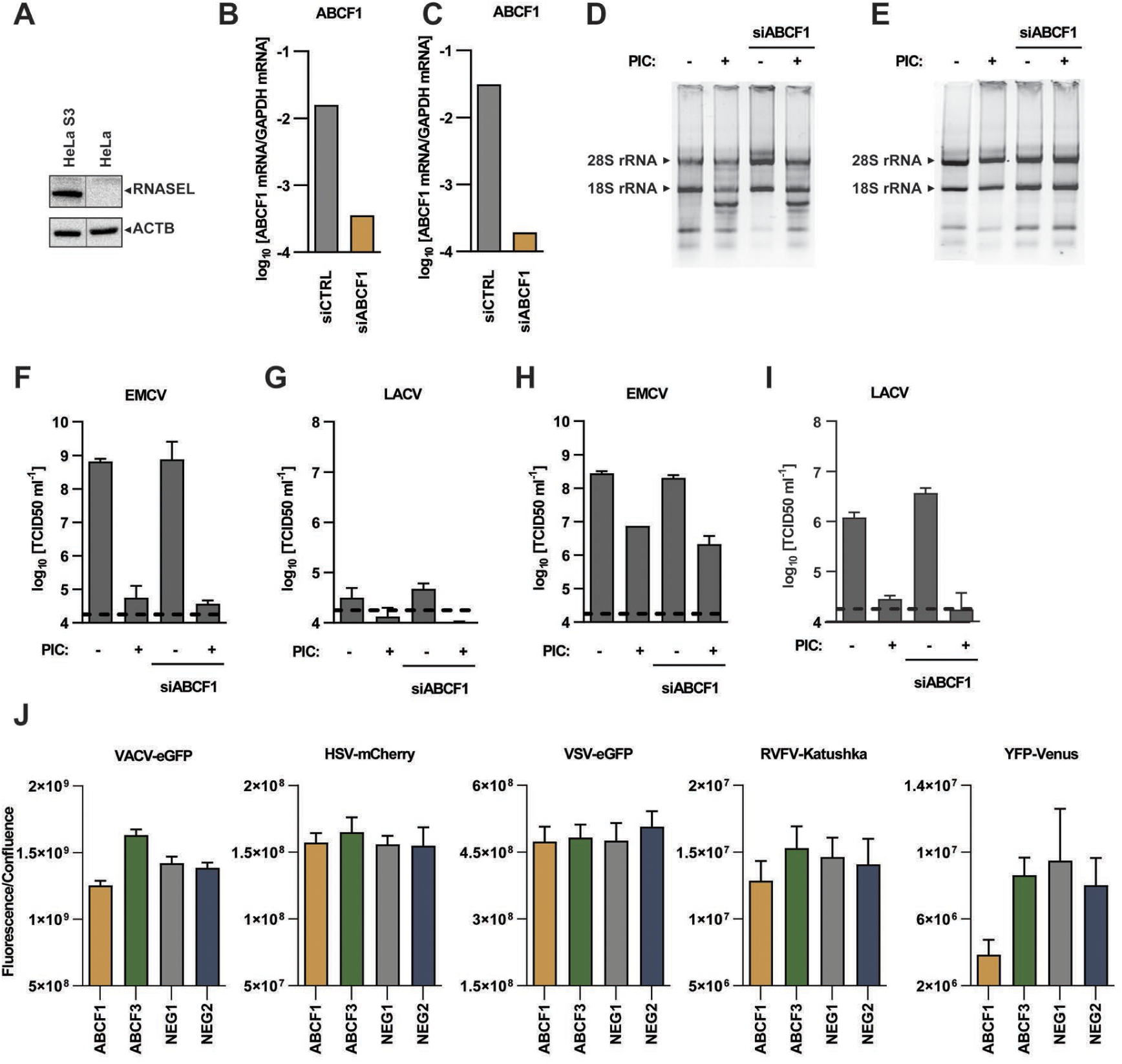
Functions of ABCF1 and RNAse L depletion in human cells. **(A)** Immunoblot analysis for endogenous RNase L in HeLa and HeLa S3 cells. **(B, C)** HeLaS3 and HeLa cells were electroporated with siRNA against ABCF1 (siABCF1) or non-targeting control siRNA (siCTRL) and expression levels of ABCF1 and GAPDH mRNA were measured by qPCR. Bar graphs show normalized average expression levels 48 h after siRNA treatment. **(D, E)** HeLa and HeLa S3 cells from (B, C) were transfected with 300ng polyI;C or mock transfected using Lipofectamine 2000. 24h later total RNA was isolated and visualized on an agarose gel. 28s and 18s rRNA are indicated. **(F, G)** HeLa S3 cells from (B) were infected with EMCV (MOI: 0.01) (F) or LACV (MOI: 1) (G) and accumulation of virus was titrated on Vero E6 cells 24h after infection. **(H, I)** as (F, G) but HeLa S3 cells were used. **(J)** A549 cells were transduced with lentiviral vectors expressing puromycin resistance, Cas9 and gRNAs targeting ABCF1, ABCF3 or non-targeting controls (NEG1 and NEG2) and selected for puromycin resistance. Obtained cells were infected with the indicated viruses expressing fluorescent reporter proteins. The bar graphs show mean fluorescent intensity (± sd, n=4) normalized to cell confluence for timepoint of 24 hpi (VSV-eGFP MOI 0.2, RVFV-Katushka MOI 0.5), 40 hpi (YFV-Venus MOI 2), 48 hpi (HSV-1-mCherry MOI 2) or 72 hpi (VACV-eGFP MOI 0.01). One representative experiment of three is shown.

### ABCF1 does not affect cellular translation and interferon-α/b induction

ABCF1 has previously been linked to efficient ^m7^G cap-dependent and EMCV IRES mediated translation (29). We envisioned that regulation of ABCF1 activity through 2′–5′ OA could affect this activity. To evaluate whether 2′–5′ OA has an effect on translation that is independent of RNase L, we employed a pulsed-SILAC mass spectrometry approach, which allows for quantification of newly synthesized proteins in a given timeframe (Figure 3a). We stimulated HeLa S3 and HeLa cells with 2′–5′ OA or dephosphorylated OH-2′–5′ OA and evaluated incorporation of heavy-labelled amino acids by LC-MS/MS within the first 12 h after stimulation (Figure 3a). As expected, compared to OH-2′–5′ OA, 2′–5′ OA transfection reduced the overall translation rates in HeLa S3 cells (Figure 3b; Supplementary Table S2), which is in line with activation of RNase L in this cell type. In contrast, the translation rates were comparable between 2′–5′ OA and OH-2′–5′ OA treated HeLa cells (Figure 3c; Supplementary Table S2), suggesting that 2′–5′ OA do not regulate translation in absence of RNase L and indicating that 2′–5′ OA interaction with alternative proteins, including ABCF proteins, does not affect overall translation.

**Figure 3:**
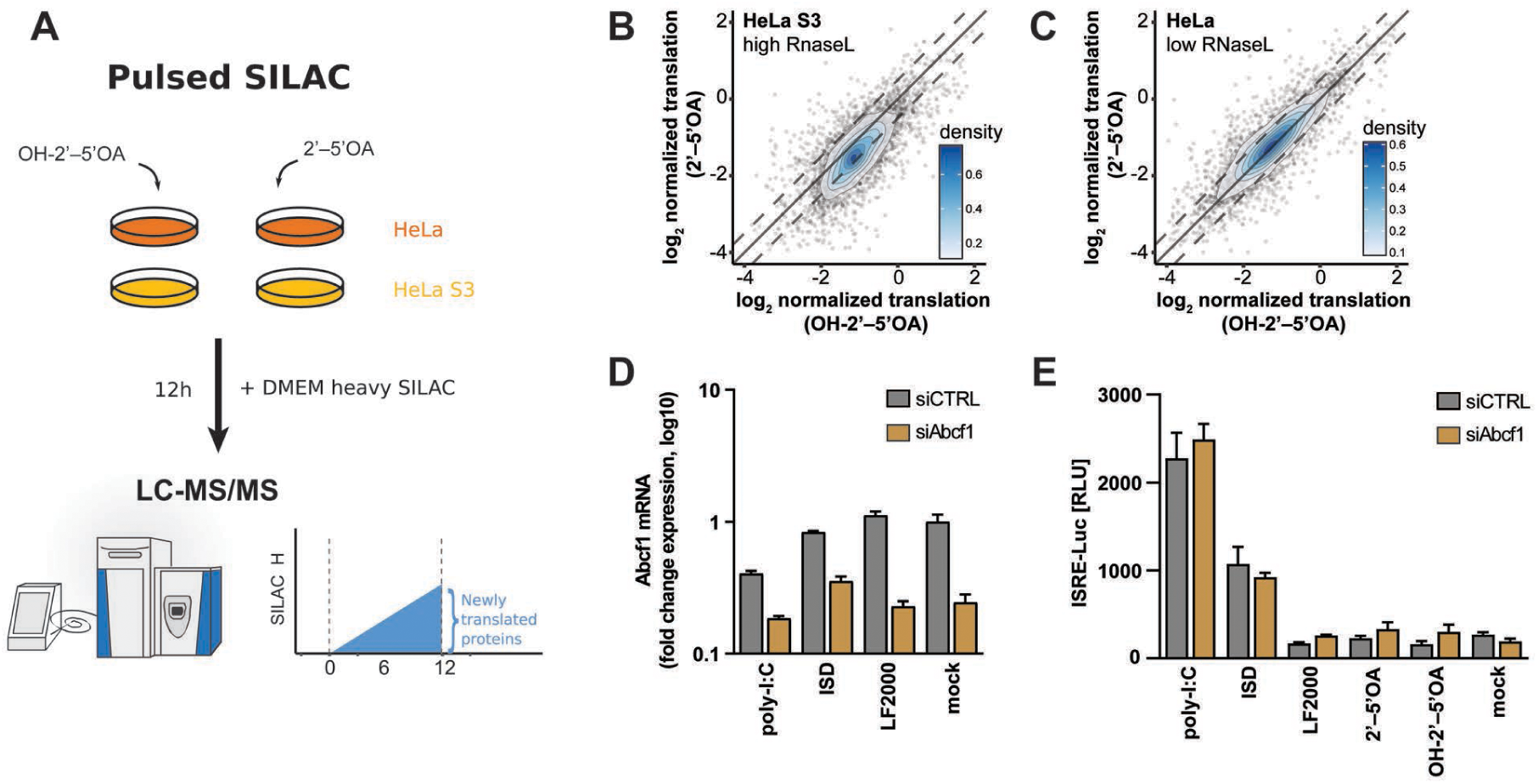
Evaluation of ABCF1 in protein translation and activation of ISRE after stimulation. **(A)** Experimental scheme of the pulsed SILAC approach used to measure protein synthesis in HeLa and HeLa S3 cells after treatment with OH-2′–5′ OA or 2′–5′ OA. **(B)** Density scatter plot comparing the normalized and log_2_ transformed intensity of heavy SILAC-labelled proteins in HeLa S3 cells treated with 2′–5′ OA or OH-of 2′–5′ OA (n = 4). Diagonal lines indicate a slope of 1, including an offset of ± log_2_ 0.5 in case of dashed lines. **(C)** As (B) in HeLa cells with low RNase L levels. **(D, E)** MEFs were treated with siRNA for ABCF1 or non-targeting control for 48 h and treated with the indicated stimuli. **(D)** Abundance of ABCF1 normalized to GAPDH transcripts in relation to mock siCTRL. Graph shows mean (± sd, n = 2) of one representative experiment of two. **(E)** MEF cells were treated with different stimuli to induce type-I interferon production. After 24 h the supernatants were assayed for presence of typ-I interferon using a bioassay on ISRE-Luc reporter containing L292 cell line. Gaph shows mean (± sd, n = 3) of one representative experiment of three.

In mouse cells, ABCF1 has been proposed to bind immunostimulatory DNA (ISD-DNA) and to regulate expression of type-I interferons (31). However, while ABCF1 bound well to 2′–5′ OA, we were unable to confirm significant binding of ABCF1 to ISD-DNA (Figure 1f). We nevertheless depleted ABCF1 in MEFs and monitored the accumulation of type-I interferon in response to poly-I:C, ISD-DNA or 2′–5′ OA (Figure 3d). Supernatants were assayed for accumulation of type-I IFN using an ISRE-luciferase expressing LL171 cell line (Figure 3e). Transfection of both, poly-I:C and ISD-DNA led to accumulation of type-I interferons. In contrast, we could not detect substantial amounts of IFN in 2′–5′ OA transfected cells. However, depletion of ABCF1 in our hands did not influence expression of type-I interferons regardless of the stimulus used (Figure 3e).

We concluded form these experiments that ABCF1 and -3 can bind to 2′–5′ OA in a phosphate dependent manner and that this binding is conserved between human, mouse and *Drosophila* cells. However, in human cells ABCF1 and -3 did not have a direct function on virus growth, RNase L activation or 2′–5′ OA dependent regulation of translation.

### 2′–5′ OA binding to Decr1 is species-specific and occurs in the NADPH co-factor binding pocket

We asked whether 2′–5′ OA may bind to proteins in a species-specific manner. Comparing mass spectrometry results between human, mouse, and *Drosophila* revealed proteins that undergo species-specific interactions with 2′–5′ OA (Figure 1 b-d). 2,4-Dienoyl-CoA reductase 1 (Decr1) was one of the highest enriched 2′–5′ OA-binding proteins identified in mouse cell lines but exhibited no enrichment from human or *Drosophila* cells. Decr1 is a mitochondria associated metabolic enzyme necessary for β-oxidation and metabolism of unsaturated CoA esters containing *trans* double bonds in both even-and odd-numbered positions and requires NADPH cofactor binding for substrate processing (32). We purified recombinant mouse Decr1 (mDecr1) and measured interaction *in vitro* using an electrophoretic mobility shift assay. We observed that mDecr1 directly binds radiolabeled 2′–5′ OA *in vitro* and forms a stable protein–RNA complex with ∼50–60 μM affinity (Figure 4a). As known for RNase L and also observed for ABCF1 (Figure 1 e-h), mDecr1 interactions with 2′–5′ OA were also diminished in the absence of a 5′ triphosphate (Figure 4a). Together these results confirm that mDecr1 directly binds to 2′–5′ OA.

**Figure 4.**
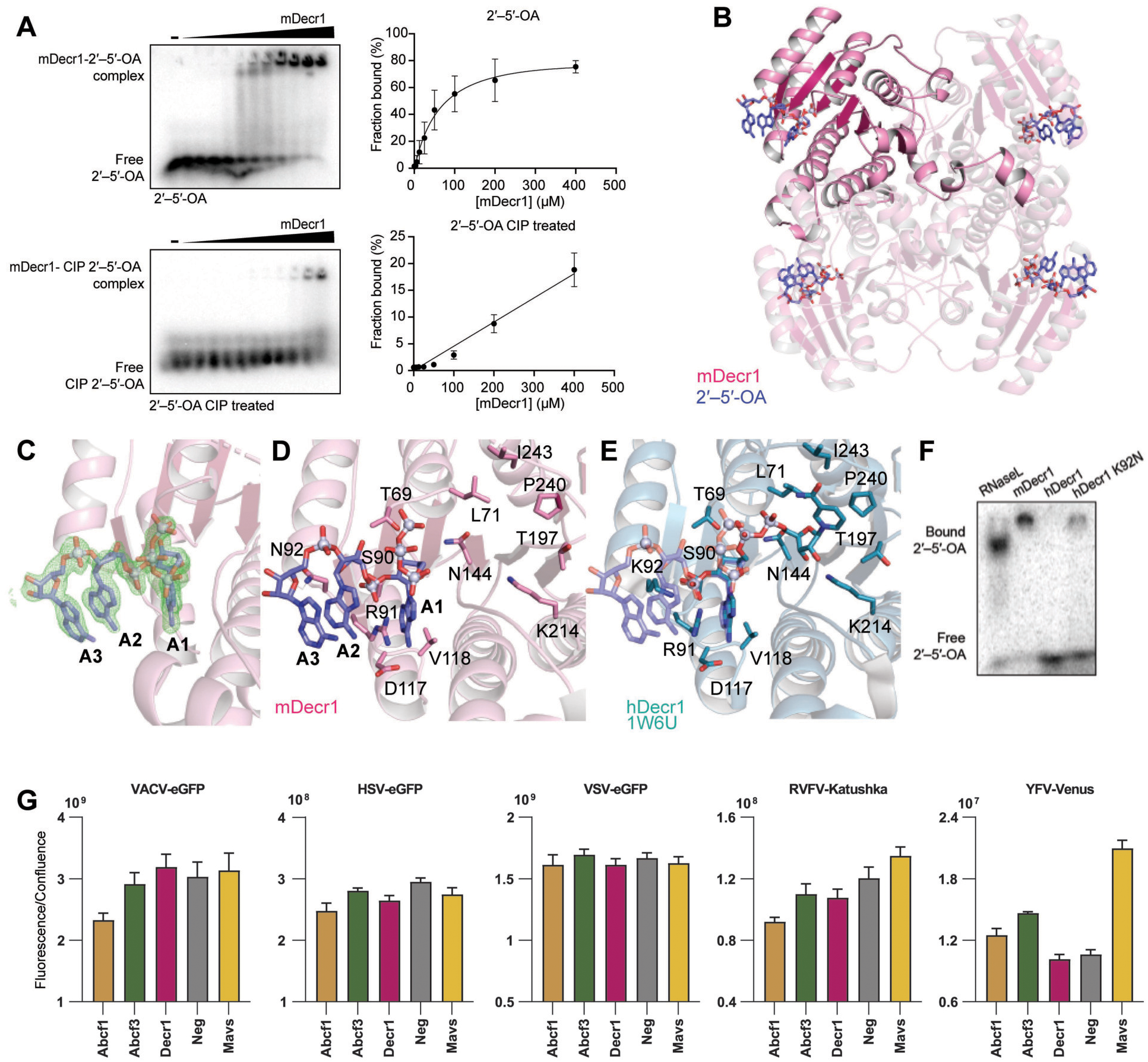
Species-specific binding of 2′–5′ OA to mDecr1. **(A)** Comparison of fraction of 2′–5′ OA bound with or without the 5′ triphosphate. Calf-intestinal phosphatase treatment removes the 5′ triphosphate and reduces mDecr1-2′–5′ OA bound complex formation. **(B)** 1.35 Å crystal structure of mouse Decr1 bound to 2′–5′ OA. Decr1 is a homo-tetrameric metabolic auxiliary enzyme that catalyzes the reduction of trans-unsaturated fatty acids in the mitochondria. We see electron density for one molecule of 2′–5′ OA with three clearly defined bases bound to each monomer of Decr1. **(C)** 2Fo-Fc map of 2′–5′ OA bound to one monomer of Decr1 contoured to 1 σ. **(D)** 2′–5′ OA bound to mouse Decr1. 2′–5′ OA contacts charged residues in each Decr1 monomer with specific contacts to the 2′ phosphate and first base. Binding pocket and active site residues in mDecr1 and hDecr1 are highly conserved (compare to E). **(E)** NADPH bound to human Decr1 (PDB: 1W6U) with 2′–5′ OA modeled and overlayed. 2′–5′ OA binds in an extended binding site compared to NADPH, which has key contacts deeper in the binding pocket. N92 in mDecr1 is permissive to 2′–5′ OA binding, while K92 in hDecr1 allows NADPH binding but sterically clashes with modelled 2′–5′ OA binding. **(F)** Comparison of mouse Decr1, human Decr1, or human mutant K92N and 2′–5′ OA complex formation. WT human Decr1 cannot form a complex with 2′–5′ OA, while the K92N mutation in the human Decr1 background is permissive to 2′–5′ OA binding and complex formation. **(G)** NIH-3T3 cells were transduced with lentiviral vectors expressing Cas9 and gRNA targeting Abcf1, Abcf3, Decr1, Mavs or non-targeting control (Neg). Targeted cell populations after puromycin selection were infected with viruses encoding fluorescent reporter genes. The graphs show mean fluorescent intensity (± sd, n = 4) normalized to cell confluence for timepoints 27 hpi (VSV-eGFP MOI 0.1), 48 hpi (RVFV-Katushka MOI 0.1), 60 hpi (YFV-Venus MOI 0.5), 72 hpi (HSV-1-eGFP MOI 1) or 96 hpi (VACV-eGFP MOI 0.01).

To explain the molecular basis of the species-specific mDecr1 2′–5′ OA recognition, we determined a 1.4 Å crystal structure of the mDecr1–2′–5′ OA complex (Figure 4b; Supplementary Table S3). mDecr1–2′–5′ OA crystals were grown in the presence of enzymatically synthesized 2′–5′ OA. Similar to previous structures of human Decr1 (hDecr1) (33), mDecr1 forms a homotetrameric enzyme with four outward facing active sites. Surprisingly, 2′–5′ OA binds directly in the mDecr1 catalytic active site with clear electron density for 2′–5′ OA observed in each protomer (Figure 4b,c). mDecr1 residues T69 and N144 coordinate the 5′ end of 2′–5′ OA making direct contact with the 5′ triphosphate and the ribose of the A1 nucleotide (Figure 4d), and by contact between D117 and the A1 adenine N1 and N6 positions. mDecr1 residues S90 and R91 coordinate the first 2′–5′ phosphodiester bond and explain specific interaction with 2′–5′-linked nucleic acid (Figure 4d), while residue N92 makes contact with both the A2 and A3 adenine nucleobases. Three bases are ordered and appear with clear electron density (Figure 4c,d). The remaining bases of the 2′–5′ OA chain extend away from the active-site, demonstrating that like RNase L, mDecR1 recognition is influenced by the first three bases of 2′–5′ OA chains (15).

We compared the mDecr1–2′–5′ OA structure with the structure of hDecr1 in complex with NADPH and hexanoyl-CoA (33). mDecr1 and hDecr1 share 85% identity at the amino acid level and residues in the catalytic pocket are highly conserved (Figure 4d,e). hDecr1 residues S90, R91 and D117 coordinate the NADPH adenine base and 2′ phosphate making near identical contacts as those observed between mDecr1 and 2′–5′ OA (Figure 4d,e). Notably, the chemical structures of NADPH and 2′–5′ OA contain identical 2′ phosphate and adenine base moieties, explaining how contacts in the Decr1 catalytic active site are able to coordinate similar interactions with both NADPH and 2′–5′ OA. In spite of the high level of conservation between hDecr1 and mDecr1, overlay of the two bound structures reveals a steric clash between hDecr1 residue K92 and the 2′–5′ OA A2 nucleobase (Figure 4e). In mDecr1, an Asn is encoded at position 92 that accommodates the extended nucleotide chain of 2′–5′ OA and enables binding (Figure 4d,e). We tested interaction between hDecr1 and 2′–5′ OA *in vitro* and confirmed that the wildtype protein is unable to form a complex with 2′–5′ OA (Figure 4f). An hDecr1 K92N substitution permits hDecr1 to weakly bind 2′–5′ OA, confirming a key role for this Decr1 residue in controlling species-specific interactions with 2′–5′ OA (Figure 4f).

Since Decr1 was binding in a species-specific manner we envisioned that potential antiviral effects of Decr1 would only be visible if tested in a murine system. The mDecr1–2′–5′ OA structure suggested that 2′–5′ OA binding would occlude the NADPH co-factor binding site and prevent enzymatic activity. Inhibition or knockdown of Decr1 leads to an accumulation of poly-unsaturated fatty acids (34, 35), and increased levels of poly-unsaturated fatty acids are associated with reduction in viral titers for enveloped viruses (36, 37). To test the effect of Decr-1, as well as the murine Abcf1 and Abcf3 on growth of different viruses we performed virus growth assays using fluorescent reporter viruses in NIH-3T3 cells that were CRISPR/Cas9 targeted for mDecr1, Abcf1, Abcf3, and controls (Neg and Mavs). Depletion of Mavs resulted in higher growth of YFV (Figure 4f), while other viruses were not affected by Mavs depletion, which is in line with their ability to perturb activation of RIG-like receptors (38). Depletion of most newly identified 2′–5′ OA binders did not result in a significantly increased growth of the viruses tested, as compared to control cells (Figure 4f,). The only exception was a modestly increased growth for yellow fever virus in Abcf3 depleted cells, which is in line with previous publications (23).

Collectively, our results clarify the species-specificity of Decr1–2′–5′ OA complex formation. Functional experiments suggest that additional factors or further assessment of cellular activities may be required to understand the biological function of Decr1 and other murine proteins in 2′–5′ OA signaling.

## Conclusions

A conserved feature of the innate immune response to viral infection is synthesis of specialized nucleotide signals that potently activate host defenses. In animal cells, OAS enzymes and cGAS-like receptors directly sense pathogen replication and then synthesize nucleotide signals including 2′–5′ OA and cGAMP that initiate antiviral immunity (2, 39–41). Likewise, the rece nt discovery of specialized nucleotide signals in prokaryotic anti-phage defense (42–44) demonstrates that nucleotide second messenger signaling is ancient and exceptionally widespread in biology.

Nucleotide second messengers may either transfer signals to a single protein and therefore serve as signal amplifier or may be able to distribute the signal to multiple proteins or even to different cells. The latter has been demonstrated for cGAMP, which has the ability to cross cell barriers through gap junctions, leading to a timely local innate immune response (45). 2′–5′ OA may similarly activate cells in the vicinity of virus infections in order to limit pathogen spread by creating a barrier of cells that do not support efficient virus replication. Here we explored the possibility that 2′–5′ OAs are binding to different target proteins within a cell and act as a signaling hub, activating a variety of different proteins without the requirement of direct protein–protein contact. In unbiased affinity proteomics analysis, we identified ABCF1, -3, E as well as murine Decr1 as specific binders of 2′–5′ OA.

The regulation of signaling events by nucleotide second messengers requires high specificity and tight control to avoid side effects such as autoimmunity or immunopathology. Most innate immune processes rely on structural and chemical features to ascertain this specificity. OAS generates oligonucleotides that bear two hallmarks not present under physiological conditions that aid in signaling discrimination: 2′–5′ linkages and a 5′ triphosphate group. Our structural data demonstrate that Decr1 binding to 2′–5′ OA requires 2′–5′ linkages and that sequence alterations in the RNA binding site of human DECR1 prevent binding of 2′–5′ OA. At this stage we do not know whether and how 2′–5′ linkages are similarly required for binding of ABCF proteins or whether such linkages would change the affinity of this interaction. However, all novel binding partners described here require triphsophorylated 5’ terminus. The 5′ triphosphate group has also been shown to be required for binding and activation of RNase L, highlighting this feature as being important to ensure signaling specificity.

We explored potential downstream effects of 2′–5′ OA binding of ABCF1, -3 and mouse Decr1. ABCF proteins have been linked to a variety of cellular effects including translational control and innate immunity. Mutations in the ATP hydrolysis sites showed reduced overall protein synthesis in HEK293 cells, as evidenced by a loss of polysomes which is a hallmark of impaired translation (29) and mutations in ABCF1 have been proposed to lead to alternative start site selection (46). However, using pulsed SILAC experiments, we could not find evidence that delivery of 2′–5′ OA regulates protein translation in an RNase L independent manner. Moreover, pulsed SILAC analysis did not reveal differences in abundance of peptides that map to 5′ N-termini of identified proteins in 2′–5′ OA transfected RNase L deficient HeLa cells (data not shown). ABCF1 has also been linked to regulation of innate immune responses after cytoplasmic delivery of DNA (31). However, unlike reported by Lee et al., in our hands ABCF1 precipitated only weakly with DNA as compared to 2′–5′ OA. Transfection of immunostimulatory DNA or 2′–5′ OA inducing poly-I:C led to similar expression of IFN-α/β in ABCF1 depleted MEFs. To establish whether ABCF proteins have antiviral activity we depleted ABCF proteins in human A549, MEFs or murine NIH3T3 cells. Collectively these data revealed an inconsistent phenotype for virus growth in ABCF1 and -3 depleted cells. It may be that additional functions that are not directly related to virus replication may be regulated or that downstream regulatory processes are only selectively available in specific cells types (47, 48). 2′–5′ OA may potentially contribute to such processes in a regulatory manner.

Decr-1 was among the proteins with the highest enrichment scores in AP-MS experiments using murine cell lysates. Our results define mouse Decr1 as novel 2′–5′ OA-binding protein and suggest auxiliary functions for OAS proteins and 2′–5′ OA signaling in animal cells (Figure 4). The micromolar binding affinity between 2′–5′ OA and mouse Decr1, compared to the nanomolar binding affinity between RNase L and 2′–5′ OA (49) may reflect its role as a secondary arm of 2′–5′ OA binding and a hierarchy of signaling outcomes beginning top down with RNase L. Some viruses encode inhibitors of RNase L that impair either RNase L function or binding to 2′–5′ OA (50) while other viruses upregulate the cellular inhibitor of RNase L, RLI, during infection (51, 52). In the event of RNase L inhibition, this secondary arm of 2′–5′ OA effects may become more pronounced and required for an alternative means of cellular metabolic shutdown.

Our results suggest that 2′–5′ OA blocks NADPH binding to Decr1 in the absence of substrate and could conceivably prevent its enzymatic activity. Knock out or inhibition of Decr1 leads to an accumulation of poly-unsaturated fatty acids (PUFAs), ER and mitochondrial stress and eventually ferroptosis (34, 35), and could provide an alternative means of cell death and metabolic shutdown during viral infection. The accumulation of lipids and PUFAs is detrimental to enveloped viruses and may hinder RNA viral replication in membrane-bound compartments (53, 54), and these cumulative antiviral effects could make up for a defective RNase L response. However, Decr-1 depletion in NIH 3T3 cells did not lead to an apparent phenotype in virus infected cells using a subset of viruses.

Notably, Decr1 is well-conserved across more ancient organisms and lower metazoans. It has not escaped our notice that the effects of this secondary arm of OAS-dependent 2′–5′ OA signaling may be indicative of more ancient pathways in organisms that do not encode RNase L, and more modern organisms may have lost these auxiliary features. As such, exploring alternative roles of 2′–5′ OA in invertebrates or lower metazoans may provide further insight into RNase L independent roles of 2′–5′ OA.

Collectively we show that 2′–5′ OA can bind to RNase L as well as additional proteins. The ability to bind 2′–5′ OA appears to be conserved in species evolution, as evidenced by the association of *Drosophila* orthologs or human ABCF1 and -3. Our results with mouse Decr1 indicate species-specific binding to 2′–5′ OA and the ongoing evolution of the system (Figure 4). Mice contain eight paralogs of OAS1, a-h, but only OAS1a and OAS1g are catalytically active (55, 56). The recent duplication and diversification of function within the OAS family of proteins in mice illustrate that the OAS innate immune axis is a mutable and rapidly evolving antiviral pathway. The role of alternative binding partners in a species-specific manner reflects a continuing pressure to adapt to diverse viral threats and suggests the parallel evolution of novel OAS functions and novel binding partners in a species-specific manner. Altogether, our identification and validation of Decr1 as an alternate binding partner of 2′–5′ OA emphasizes the possibility of diverse antiviral pathways controlled by a diffusible second messenger and highlights areas for future focus.

## Acknowledgements

We want to acknowledge the immunopathology of virus infections laboratory for critical discussions and suggestions. We further thank Simon Giosele, Leopold Urich, Sabine Suppman and Stephan Uebel, for support, Christopher G. Proud, Bob Silverman, Giulio Superti-Furga, Georg Kochs and Andrew Bowie for critical reagents. Work in the author’s laboratories was supported by an ERC consolidator grant (ERC-CoG ProDAP, 817798), the German Research Foundation (PI 1084/3, PI 1084/4, PI 1084/5 and TRR179, TRR237) and the Bavarian State Ministry of Science and Arts (Bavarian Research Network FOR-COVID) to AP. The work was funded in part by the Richard and Susan Smith Family Foundation and the Parker Institute for Cancer Immunotherapy (P.J.K.). A.A.G. is supported by a United States National Science Foundation Graduate Research Fellowship.

## Author Contributions

A.A.G., A.W.B., C.U., M.H. and R.H. conducted experiments. A.A.G., P.J.K., C.U. and M.H. analyzed data. A.A.G., A.W.B., P.J.K and A.P. designed the experiments and wrote the paper.

## Data Availability

The mass spectrometry proteomics data have been deposited to the ProteomeXchange Consortium (http://proteomecentral.proteomexchange.org) via the PRIDE partner repository with the dataset identifier PXD030134.

## METHODS

### Cells, plasmids and reagents

HeLa S3 (CCL-2.2 ATCC), THP-1 (300356 – CLS), Vero E6 cells (CRL-1586) all ATCC, HeLa cells were a kind gift of Giulio Superti-Furga (CeMM, Vienna, Austria), MEFs were a kind gift of Bob Silverman (Cleveland Clinic, Cleaveland, OH, USA), NIH-3T3 and LL171 reporter cells were from Georg Kochs (Uniklinik Freiburg, Germany), iBMDM were a gift from Andrew Bowie(Trinity College Dublin, Dublin, Ireland), Schneider S2 cells were from Irene Ferreira (MPI of Biochemistry, Munich, Germany). Mammalian cell lines were maintained in DMEM (Sigma) or RPMI (Sigma) for THP-1 cells, supplemented with 10% FCS (GE Healthcare) and antibiotics (100 U ml^−1^ penicillin and 100 µg ml^−1^ streptomycin). Schneider S2 cells were maintained in Schneider’s medium (Biowest) supplemented with 10% fetal calf serum, Glutamax (Invitrogen) and antibiotics (100 U ml^−1^ penicillin, 100 μg ml^−1^ streptomycin).

The plasmids pCMV5-HA-hABCF1, pCMV5-HA-hABCF1-E439Q/E730Q and pCMV5-HA-hABCF1-K304M/K626M were a kind gift from Christopher G. Proud (University of Dundee/ University of Adelaide) and were used to create pETG10A-hABCF1, pETG10-hABCF1_A-/- and pETG10-hABCF1_B-/- and pCoofy4-hABCF1 to produce recombinant human ABCF1 in *E. coli*. Antibodies against the following proteins were used: ABCF1 (Aviva Systems biology; ARP43631_P050), ABCF3 (Sigma; HPA036332), RNASE L (Abcam; ab13825), HA tag (HRP coupled, Sigma; H6533), β-actin (HRP coupled, Santa Cruz; sc-47778) and HRP coupled secondary antibodies against rabbit IgG (Cell Signaling Technology; 7074) and against murine IgG (Columbia Biosciences; HRP-112).

### Treatments, transfections, knockdowns and generation of stable cells lines

Generation of phosphorylated 2′–5′ OA is described below (Aarhus University); for dephosphorylation they were treated with Alkaline Phosphatase Calf Intestinal (New England Biolabs). Poly-I:C (Sigma-Aldrich, P9582), ISD-45, 2′–5′ OA and dephosphorylated OH-2′–5′ OA were transfected into cells using Lipofectamine 2000 (Invitrogen) according to the protocol provided by the manufacturer.

For siRNA-mediated knock-down, siRNA (0.4 nmol ml^−1^ Hela, HeLa S3 or MEFs) was electroporated using the Neon Transfection System (Invitrogen) according to cell line specific instructions of the manufacturer. The following siRNA were used: Control: AAGGTAATTGCGCGTGCAACT Core Facility of Max Planck Institute of Biochemistry (Martinsried, Germany), Pooled human ABCF1: (#1: AAGGGAAGGCTAAGCCTCAAA, #2: CAGAGTGTTAGCCAAATCGAT, #3: CTGGCTTAATAACTACCTCCA, #4: CCCAGCGGCTCCACTACTATA) (Qiagen), Pooled murine ABCF1 (#1: AGGAAGUCCUGACUCGAAA, #2: CGAUGAUAGUGAUGAGAGA) Core Facility of Max Planck Institute of Biochemistry (Martinsried, Germany). siRNA Knockdown efficiency was tested as follows: 200 ng of RNA was reverse transcribed with Prime Script RT Master Mix (Takara) and quantified using QuantiFast SYBR Green RT-PCR Kit (Qiagen) and a CFX96 Touch Real-Time PCR Detection thermocycler (BioRad). The following primers were used for quantitative RT-PCR: human ABCF1 (5′ AGAGCACGAGCCCATCAG 3′, 5′ TCTTCCCCAGCCTGTTTATC 3′), human GAPDH (5′ GATTCCACCCATGGCAAATTC 3′, 5′ AGCATCGCCCCACTTGATT 3′), mouse TBP (5′ CCTTCACCAATGACTCCTATGAC 3′, 5′ CAAGTTTACAGCCAAGATTCA 3′), mouse Abcf1 (5′ AGAAAGCCCGAGTTGTGTTTG 3′, 5′ GCCCCCTTGTAGTCGTTGATG 3′)

To generate knockout cells CRISPR/Cas9 technology was used. gRNA sequences of target genes were cloned into pLentiCRISPR-v2 (Addgene #52961) and lentiviral vectors encoding gRNA and Cas9 were generated in 293T. Briefly, psPAX2 (Addgene #12260) and pMD2.G (Addgene #12259) and gRNA containing plasmid lentiCRISPRv2 were transfected into 293T cells using polyethylenimine (Polysciences). Supernatants were collected at 48 and 72 hours post transfection, filtered and stored at –80°C. Knockout cell lines were obtained by transduction in the presence of polybrene (8 μg ml^−1^) followed by antibiotic selection with 2 μg ml^−1^ puromycin for 4 passages. Following gRNA sequences were used against mouse and human genes:

Abcf1 (mouse): CCGATCCACTCGGGCTCGGG; GGCGGCCGGCTACTATGTAC; AGGGCCAACCGACTTACTCC, Abcf3 (mouse) GCGAGTTCCCTGAAATTGAC; ACGGCTAGACGGTTACCGGG; GGAAAGTGACACCGTACGAG, Decr1 (mouse) ATCCGGATGTAAGTGTAACG; TCTGAGGACCCGGATCGCAG; AGTCGGGTCCAAACGGCTAA, Mavs (mouse) GCCACCAGACATCCTCGCGA; CCAACTCCGGGGCCGTCGCG; TAATTTTGATGGCGCTGTAT, Neg (mouse) AAGCGGTGGGTGTCGATAAT; CTTCCTAGCCATAGCCGCGT,

ABCF1 (human) GCAACACATCAATGTTGGGA; TAAGCCAGATGACAGCGTTG; TGTAATTGCCCCTATAGTAG, ABCF3 (human) CAGCGGCTAGATGGTTACCG; TGCGAGAGGATTTGCTACGG; GCAGAGTGTTGTACATGCGC, NEG1 (human) TGCAAAGTTCAGGGTAATGG; AATAACGCGTAACTCCCACC; AGCTTGACAATGCACACTAC, NEG2 (human) ACCGGAAACGATGAGTGGGG; AAGCGGTGGGTGTCGATAAT; CTTCCTAGCCATAGCCGCGT

### Recombinant protein production

Human recombinant ABCF1 with N-terminal 6×His-MBP tag was obtained in *E. coli* expression system (expression vector pCoofy4) and purified by Core Facility of Max Planck Institute of Biochemistry (Martinsried, Germany) (Anna s paper). Recombinant human OAS1, human Decr1, human RNase L, and mouse Decr1 were cloned from synthetic DNA (IDT) into a custom pET vector containing a 6×His-SUMO2 solubility tag. Tagged protein was expressed in E. coli strain BL21-RIL (Agilent) containing the rare tRNA plasmid pRARE2 as previously described (57, 58). Transformed colonies were grown in a 30 ml MDG media starter culture (0.5% glucose, 25 mM Na_2_PO_4_, 25 mM KH_2_PO_4_, 50 mM NH_4_Cl, 5 mM Na_2_SO_4_, 2 mM MgSO_4_, 0.25% aspartic acid, 100 mg ml^−1^ ampicillin, 34 mg ml^−1^ chloramphenicol, and trace metals) overnight at 37°C, and used to inoculate 1 L M9ZB media cultures (0.5% glycerol, 1% CAS amino acids, 47.8 mM Na_2_PO_4_, 22 mM KH_2_PO_4_, 18.7 mM NH4Cl, 85.6 mM NaCl, 2 mM MgSO_4_, 100 mg ml^−1^ ampicillin, 34 mg ml^−1^ chloramphenicol, and trace metals) at OD600 ∼0.05. Cultures were grown at 37°C, 230 RPM until OD600 ∼2.5, chilled on ice for 10 min, induced with 0.5 mM IPTG, and incubated at 16°C for shaking at 230 RPM for ∼16 h before harvest. Cultures were pelleted, lysed by sonication in 1× Lysis Buffer (20 mM HEPES-KOH pH 7.5, 400 mM NaCl, 30 mM imidazole, 1 mM DTT and 10% glycerol). Protein was purified from sonicated lysates using Ni-NTA resin (Qiagen) and gravity flow. Ni-NTA resin was washed with 1× Lysis Buffer supplemented to 1 M NaCl and eluted with 1× Lysis Buffer supplemented to a final concentration of 300 mM imidazole. For crystallography, purified protein was treated with recombinant human SENP2 protease (D364–L589, M497A) to remove the SUMO2 tag and dialyzed for ∼16 h against Dialysis Buffer containing 20 mM HEPES-KOH pH 7.5, 250 mM KCl, and 1 mM DTT. The next morning, protein was concentrated using a 30K-cutoff concentrator (Millipore) and purified by size exclusion chromatography (SEC) using a 16/600 Superdex 200 column with Gel Filtration Buffer containing 20 mM HEPES-KOH pH 7.5, 250 mM KCl, 1 mM TCEP. Purified SEC fractions were pooled and concentrated to >30 mg ml^−1^, aliquoted in 50 μl, flash frozen in liquid nitrogen, and stored at −80°C. Protein for biochemical assays was purified similarly except the human SENP2 protease step was omitted, resulting in tagged protein.

### Virus infections, measurement of virus growth and IFN bioassay

The following viruses were used in the study: Encephalomyocarditis virus (EMCV), La Crosse encephalitis virus (LACV rec wt), herpes simplex virus 1 (HSV1(17+)Lox-mCherry), Rift Valley fever virus Clone 13 (RVFV-Katushka), vaccinia virus (VACV-GFP), yellow fever virus (YFV-Venus), vesicular stomatitis virus (VSV-GFP). Supernatants from EMCV and LACV infected cells (MOI = 0.01 and 1 respectively) were collected at 24 hpi and titrated on the Vero E6 cells using the TCID50 method. Knock out A549 and NIH-3T3 cell lines were infected with viruses encoding fluorescent reporters at indicated MOI and the reporter expression was measured using Incucyte S3 automated fluorescence light microscope (Essen Bioscience) every 3 h. The phase image and the fluorescent signal in red or green channel were collected. The background was removed using the built-in analysis module (IncuCyte S3 Software, Essen Bioscience, Version: 2020C Rev1). Integrated intensity of fluorescent signal was normalized to confluence for each picture.

Mouse type-I interferon secreted by MEFs was measured using the interferon bioassay. Collected MEF supernatants were added to the LL171 cells (containing ISRE-firefly luciferase reporter). After 16 h cells were lysed in 1x passive lysis buffer (Promega) and firefly luciferase activity was measured after addition of firefly substrate-containing buffer (20 mM Tricine, 3.74 mM MgSO4, 33.3 mM DTT, 0.1 mM EDTA, 270 μM Coenzyme A trilithium salt (Sigma-Aldrich #C3019), 470 μM D-Luciferin sodium salt (Sigma-Aldrich #L6882), 530 μM ATP disodium salt (Sigma-Aldrich #A7699), pH 7.8-8).

### Affinity purification, mass spectrometry experiments and bioinformatic analyses

Biotinylated 2′–5′ OA were synthesized according to the protocol described in Turpaev et al. (59) with minor modifications. 10 µg porcine OAS1 isoform was produced and purified as described in (60) and mixed with 2 mM ATP and 25 µg of poly-I:C, in 1 ml of a buffer containing 20 mM TRIS, 0.1 mM EDTA, 10 mM MgCl2 and 0.2 mM DTT, Ph 7.8 (all concentrations are final concentrations), incubated at 37°C for 2 h, loaded onto a 5 ml Source Q column and eluted using the gradient described in (59). To biotinylate 2′–5′ OA the same reaction was performed with purified trimeric 2′–5′ OA and biotinylated dATP as substrates. Biotinylated ISD-45 was prepared by annealing of biotin-labeled oligonucleotides. For the enrichment of proteins using a biotinylated baits, 60 µl streptavidin affinity resin (Novagen) was first incubated either with 300 pmol 2′–5′ OA or 300 pmol ATP in TAP buffer (50 mM Tris pH 7.5, 100 mM NaCl, 5% (v/v) glycerol, 0.2% (v/v) Nonidet-P40, 1.5 mM MgCl2 and protease inhibitor cocktail (cOmplete, EDTA-free; Roche), in the presence of 40 U RNase inhibitor (Fermentas) for 1 h at 4°C on a rotary wheel. Beads were washed three times with TAP buffer to remove excess of unbound nucleic acids. Cell lysates were prepared by flash-freezing cells in liquid nitrogen, followed by lysis in TAP buffer for 30 min on ice. Lysates were clarified by centrifugation at 16,000 × g for 10 min and the protein concentration was determined using a Lowry assay (DC Protein Assay, BioRAD). Nucleic acid coated beads were incubated with 2 mg protein of cell lysates for 60 min and then washed three times with TAP buffer. For WB analysis after affinity purification (lysates from THP, and HeLa cells), 5× Laemmli buffer (250 mM Tris HCl, 100 mM DTT, 50% glycerol, 10% SDS, 0.1% Bromophenol Blue) was added to the beads after washing and the samples were heated (10 min at 95°C) and analyzed by SDS-PAGE followed by immunoblotting analysis (PVDF). Affinity purification coupled to mass spectrometry (AP-MS) was performed on lysates from HeLa, immortalized murine BMDM, and drosophila S2 cells. After affinity purification and three washings with TAP buffer, the beads were additionally washed twice with TAP buffer lacking Nonidet-P40 to remove residual detergent. At least three independent affinity purifications were performed for each bait (biotinylated 2′–5′ OA or biotinylated ATP, which was used as control). Enriched proteins were denatured in U/T buffer (6 M urea, 2 M thiourea, 1 mM DTT (Sigma), 10 mM HEPES, pH 8) for 30 min and alkylated with 5.5 mM iodoacetamide (Sigma) for 20 min. After digestion through addition of 1 µg LysC (WAKO Chemicals USA) at room temperature for 3 h, the suspension was diluted in 50 mM ammonium bicarbonate buffer (pH 8). Beads were removed by filtration through 96-well multiscreen filter plates (Millipore, MSBVN1210) and the protein solution was digested with 0.5 µg trypsin (Promega) overnight at room temperature. Peptides were purified on StageTips with three C18 Empore filter discs (3M) and analyzed by MS as described previously (61). Briefly, peptides were eluted from StageTips and separated on a C18 reversed-phase column (Reprosil-Pur 120 C18-AQ, 3 µM, 150×0.075 mm; Dr. Maisch) by applying a 5% to 30% acetonitrile gradient in 0.5% acetic acid at a flow rate of 250 nl min^−1^ with a total length of 120 or 130 min, using an EASY-nanoLC system (Proxeon Biosystems). The nanoLC system was directly coupled to the electrospray ion source of an LTQ-Orbitrap XL mass spectrometer (Thermo Fisher Scientific) operated in a data dependent acquisition mode with a full scan (300 – 1,650 m/z) in the Orbitrap cell at a resolution of 60,000 and concomitant isolation and fragmentation of the ten most abundant precursor ions.

To measure translation rates by pulsed SILAC, HeLa and HeLa S3 cells were cultured in DMEM medium, containing antibiotics, 10 mM L-glutamine, 10% dialyzed fetal calf serum (PAA Laboratories) and either heavy (84 mg/l ^13^C_6_ ^15^N_4_ L-arginine and 146 mg/l ^13^C_6_ ^15^N_2_ L-lysine) or medium (84 mg L^−1^ ^13^C_6_ L-arginine and 146 mg L^−1^ ^2^H_4_ L-lysine) SILAC amino acids (Cambridge Isotope Laboratories). Medium SILAC labeled HeLa or HeLa S3 cells were used as spike-in control. These cells were labeled for at least 8 doublings with medium SILAC medium. For pulsed SILAC, HeLa and HeLa S3 cells were stimulated through MetafectenePRO-based transfection of 200 pmol 2′–5′ OA or OH-2′–5′ OA and incubated with heavy L-lysine and L-arginine amino acids for 12 h post stimulation using heavy SILAC medium. Cells from the pulsed SILAC experiment as well as medium SILAC-labelled spike-in cells were lysed in SDS lysis buffer (50 mM Tris pH 7.5, 4% sodium dodecyl sulfate) at 95°C for 5 min, sonicated for 15 min with a Bioruptor (Diagenode) and centrifuged for 5 min at 16,000 × g at room temperature. Protein concentration was determined using a Lowry assay (DC Protein Assay, BioRAD) and each sample from the pulsed SILAC experiment was mixed in a 1:1 ratio with medium SILAC-labelled proteins from the spike-in control. Subsequently, 50 µg protein per sample were reduced with 10 mM DTT (Sigma) for 30 min, alkylated with 55 mM iodoacetamide (Sigma) for 20 min at room temperature, and precipitated with 80% acetone for 3 h at −20°C. After centrifugation for 15 min at 16,000 × g at 4°C, protein pellets were washed with 80% acetone, dried for 30 min at room temperature and dissolved in U/T buffer (6 M urea, 2 M thiourea, 1 mM DTT, 10 mM HEPES, pH 8). Proteins were digested with LysC and trypsin overnight at room temperature and purified on StageTips with 3 layers of C18 Empore filter discs (3M) and analyzed by LC-MS/MS using an Easy nanoLC system coupled to a Q-Exactive HF mass spectrometer (Thermo Fisher Scientific). Peptide separation was achieved on a C18-reversed phase column (Reprosil-Pur 120 C18-AQ, 1.9 µM, 500×0.075 mm; Dr. Maisch) using a linear gradient of 2% to 30% acetonitrile in 0.1% acetic acid with a total length of 180 min. The mass spectrometer was set up to run a Top15 method, with a full scan scan range: 300-1,650 m/z, R: 120,000, AGC target: 3e6, max IT: 20 ms) followed by isolation, HCD fragmentation and detection of the 15 most abundant precursor ions (scan range: 200-2,000 m/z, R: 15,000, AGC target: 1e5, max IT: 25 ms, NCE: 27, isolation window: 1.4 m/z, dynamic exclusion: 20 s).

RAW mass spectrometry files were processed with MaxQuant (version 2.0.1.0) (62) using the built-in Andromeda search engine and protein sequence data from either human, mouse or fly proteomes (UniprotKB, release 2021-06). In addition to the standard settings, iBAQ and label free quantification (LFQ) was enabled with an LFQ minimum ratio count of 1, disabled LFQ normalization and without stabilization of large LFQ ratios. Match between runs was activated but only allowed across replicates of the same condition. In case of SILAC data, the multiplicity was set to 3 to include light (Lys0/Arg0), medium (Lys4/Arg6) and heavy (Lys8/Arg10) SILAC amino acids with a maximum number of 3 labeled amino acids per peptide. Data from the proteinGroups output table of MaxQuant was used for all downstream statistical analyses using Perseus (version 1.6.15.0) (63), R (version 4.1.0) and RStudio (version 1.4.1717). All reverse sequence hits, potential contaminants and proteins which were only identified by site were removed. In case of AP-MS experiments, LFQ intensities were log_2_-transformed and normalized by subtracting the sample-specific median intensity. Proteins were further removed if they were not quantified in at least 2 replicates in either of the conditions and missing values were imputed for each replicate individually by sampling values from a normal distribution calculated from the original data distribution (width = 0.3 × standard deviation, downshift = −1.8 × standard deviation). Differentially enriched proteins between the 2′–5′ OA versus ATP affinity purifications were identified via two-sided Student’s t-tests (S0 = 0) corrected for multiple hypothesis testing applying a permutation-based false discovery rate (FDR < 0.01, 250 randomizations). Significantly enriched proteins with a log2 fold change of ≥ 1.5 were considered as hits.

For analysis of the pulsed SILAC data, SILAC channel-specific light (L, degradation), medium (M, spike-in control) and heavy (H, translation) iBAQ intensities were used. Following log2-transformation, L and H intensities for each protein and sample were normalized by subtracting the corresponding M intensity to get normalized L/M and H/M intensity ratios. The condition-specific median L/M and H/M intensity ratios for each protein were calculated and used for visualization.

### Crystallography and structure determination

Recombinant purified mouse Decr1 was crystallized in a hanging-drop format at 18°C. Concentrated mouse Decr1 protein stocks were diluted to 10 mg ml^−1^ in buffer (20 mM HEPES-KOH, 10 mM KCl, 1 mM TCEP-KOH) and incubated on ice with ∼2 mM enzymatically synthesized 2′–5′ OA for 20 min before setting trays. Optimized crystals were grown in EasyXtal 15-well hanging-drop trays (NexTal) in 2 μl drops mixed 1:1 over a 350 μl reservoir solution of 34% PEG-300, 100 mM Tris_HCl pH 8.6). Crystals were grown for 24 h, cryoprotected using reservoir solution supplemented with 10% ethylene glycol and ∼2 mM 2′–5′ OA, and harvested by freezing in liquid nitrogen.

X-ray data were collected at the Lawrence Berkeley National Laboratory Advanced Light Source beamline 8.2.2 (DE-AC02-05CH11231) supported in part by the ALS-ENABLE program (P30 GM124169-01), and at the Northeastern Collaborative Access Team beamline 24-ID-E (P30 GM124165), and used a Pilatus detector (S10RR029205), an Eiger detector (S10OD021527) and the Argonne National Laboratory Advanced Photon Source (DE-AC02-06CH11357). X-ray data were processed using XDS and AIMLESS with the SSRL autoxds script (A. Gonzalez, Stanford SSRL). Data were processed using PHENIX (64) and the mouse Decr1 structure was determined using molecular replacement with the human Decr1 structure as a model (PDB: 1W6U) (33). Model building was performed in Coot (65), and refinement was performed in PHENIX.

### Affinity measurements by MST and Electrophoretic mobility shift assays

For microscale thermophoresis based affinity measurements the serial dilutions of human recombinant ABCF1 (15 nM – 7.16 μM) were mixed 1:1 with fluorescently labeled oligos (100 nM Alexa-488-labelled 2′–5′ OA and OH-2′–5′ OA) prepared in 1xMST buffer (50 mM Tris HCl; 150 mM NaCl; 10 mM MgCl2; 0.05% Tween-20) and measured in Monolith NT.115 (Nano Temper Tech). The instrument was set at 40% power of blue LED and 60% MST power with the on time 30 s and off time of 5 s. The normalized fluorescence was plotted against the concertation of the recombinant protein and KD calculated.

Electrophoretic mobility shift assays: Recombinant protein and enzymatically synthesized, radiolabeled 2′–5′ OA were incubated in vitro. 10 μl reactions in binding buffer containing 10 mM Tris-HCl pH 7.5, 50 mM KCl, and 1 mM DTT and 1 μM α^32^P-radiolabeled 2′– 5′ OA and either 400 μM mouse Decr1 or human Decr1, or serial dilutions (400 nM, 700 nM, 1 μM, 3 μM, 6.25 μM, 12.5 μM, 25 μM, 50 μM,100 μM, 200 μM, 400 μM) of mouse Decr1. Reactions were incubated at 25°C for 30 min prior to resolution on a 7.2 cm, 6% nondenaturing polyacrylamide gel run at 100 V for 30 min in 0.5× TBE buffer. The gel was dried at 80°C for 1 h prior to exposure to a phosphor-screen for ∼5 h to overnight and imaged using a Typhoon Trio Variable Mode Imager (GE Healthcare).

Quantification of mouse Decr1-2′–5′ OA complex formation was carried out using ImageQuant TL v8.2.0.0 software. Pixel intensity for each mouse Decr1-2′–5′ OA complex and total pixel intensity for each lane was calculated, and percent binding was determined by dividing pixel intensity for each mouse Decr1-2′–5′ OA complex by total pixel intensity for each lane. Data were fit using a non-linear regression curve for normalized one-site binding in GraphPad Prism version 9.0.0.

