## Supplementary Table 1 for "RNase L activating 2′–5′ oligoadenylates bind ABCF1, -3 and Decr-1"

**Table 1. Summary of Crystallographic Statistics**

| | *Mus musculus* Decr1–2′–5′ OA |
|---|---|
| **Data Collection** | |
| Resolution (Å)[a] | 67.39–1.35 (1.37–1.35) |
| Wavelength (Å) | 0.97918 |
| Space group | $C\ 2\ 2\ 2_1$ |
| Unit cell: a, b, c (Å) | 82.38 117.19 232.25 |
| Unit cell: α, β, γ (°) | 90.00 90.00 90.00 |
| Molecules per ASU | 4 |
| Total reflections | 1669847 |
| Unique reflections | 244100 |
| Completeness (%)[a] | 99.8 (97.6) |
| Multiplicity[a] | 6.8 (5.7) |
| $I/\sigma I$[a] | 10.6 (1.2) |
| CC(1/2)[b] (%)[a] | 99.8 (52.5) |
| Rpim[c] (%)[a] | 3.1 (55.9) |
| Sites | |
| **Refinement** | |
| Resolution (Å) | 67.39 – 1.35 |
| Free reflections | 2000 |
| R-factor / R-free | 15.1 / 16.0 |
| Bond distance (RMS Å) | 0.008 |
| Bond angles (RMS °) | 1.084 |
| **Structure/Stereochemistry** | |
| No. atoms: protein | 8490 |
| No. atoms: ligand | 436 |
| No. atoms: water | 954 |
| Average B-factor: protein | 23.82 |
| Average B-factor: ligand | 36.02 |
| Average B-factor: water | 34.82 |
| Ramachandran plot: favored | 97.55% |
| Ramachandran plot: allowed | 2.45% |
| Ramachandran plot: outliers | 0.00% |
| Rotamer outliers | 1.20% |
| MolProbity[d] score | 1.36 |
| Protein Data Bank ID | **7UCW** |

[a] Highest resolution shell values in parentheses
[b] (Karplus and Diederichs 2012)
[c] (Weiss 2001)
[d] (V. B. Chen et al. 2010)
